# NK cell immunotherapy administered at the time of HIV recrudescence is associated with viral control

**DOI:** 10.1101/2025.11.19.688897

**Authors:** Liliana K. Thron, Pongthorn Pumtang-on, Jae Woong Chang, Maxwell E. Cantor, Ian Gorrell-Brown, Kelsie L. Becklin, Zoe E. Quinn, Ahmad F. Karim, Aaron K. Rendahl, Mary S. Pampusch, Sofia A. Casares, Vaiva Vezys, Branden S. Moriarity, Pamela J. Skinner

## Abstract

One barrier to developing an HIV-1 cure is viral reservoirs persisting within B cell follicles of lymphatic tissues, partly due to failure of HIV-specific cytotoxic cells to express the follicular-homing receptor CXCR5. We evaluated HIV-specific CAR/CXCR5 NK cells armored with IL-15 and a PD-1 knockout for *in vitro* functionality. The safety and efficacy of these NK cells were assessed *in vivo* by treating ART-suppressed, HIV-infected-humanized-DRAGA mice following ART interruption. *In vitro,* CAR and control NK cells proliferated and killed HIV-Envelope-expressing cells. *In vivo*, there were no adverse health outcomes associated with cell infusions. CAR NK-treated animals had, on average, 1.88 times higher NK cell levels, and cells persisted up to 2 weeks longer than control NK-treated animals. At 56 days post-treatment, 62.5% of CAR NK-treated and 50% of control NK-treated groups had viral loads below the detection limit compared to 0% of the saline control group. Importantly, animals that had undetectable viral loads at 56 days post-treatment had earlier viral rebound post-ART interruption that coincided with high levels of NK cells, suggesting that timing of treatment with viral recrudescence was key to efficacy. This study demonstrates the promise of NK therapies for treating HIV infections.

## Introduction

Over 39.9 million individuals are living with HIV-1 worldwide ^1^. Development of efficacious therapeutics is crucial for public health and for the operational readiness of service members. Although antiretroviral therapy (ART) has substantially reduced HIV-1-related morbidity, it is costly, requires lifelong adherence, and non-adherence can lead to the emergence of ART-resistant HIV. Thus, innovative strategies to induce lifelong viral suppression are crucial. A major barrier to developing such strategies is the persistence of HIV viral replication in lymphoid B cell follicles ^2–8^, even in the presence of ART ^9–13^. The majority of HIV-specific cytotoxic cells are located outside of follicles ^14–19^, indicating that low levels of cytotoxic cell accumulation in follicles permit ongoing viral replication. Most virus-specific cytotoxic cells within lymphatic tissues fail to express the chemokine receptor CXCR5 ^14^, a follicular homing molecule that induces migration to CXCL13 in follicles. The paucity of CXCR5+ cytotoxic cells leads to failure to accumulate within B cell follicles. These findings support the development of HIV-specific immune cell therapies that express CXCR5 to enable homing to follicles and eliminate virus replication.

Chimeric Antigen Receptor (CAR) T cells have been effective in treating various cancers ^20–23^, and are also being explored to treat HIV ^24–27^. A second-generation bispecific CAR has been developed containing the HIV binding domain of CD4 linked to the carbohydrate recognition domain of mannose-binding lectin (MBL), both with high specificity for the HIV Envelope (Env) glycoprotein ^28^. The HIV-Env targeting CD4-MBL CAR shows high specificity and efficacy ^28,29^. We previously engineered therapeutic T cells to express a simian version of this CAR and CXCR5^29,30^. CAR/CXCR5 T cell-treated rhesus macaques maintained lower viral loads and follicular viral RNA levels compared to untreated control animals for the duration of the study ^29,31^. The findings suggest that CAR/CXCR5 T cells may lead to long-term viral suppression despite only persisting for 2-4 weeks, perhaps by allowing the host memory virus-specific-T cell response time to accumulate at sufficient levels to suppress remaining viral replication.

As an alternative to T cells, NK cell therapies are attractive for many reasons ^32^. To name a few: peripheral blood (PB)-derived NK cells are easy to isolate and can be expanded to clinically relevant numbers using feeder cells expressing membrane-bound IL21 (mbIL21) and 4-1BBL (Clone 9 K562s) ^33–37^. PB-NK cells do not express CD4, meaning they are not susceptible to HIV infection ^38,39^, and they do not induce graft-versus-host disease, allowing them to be used as an allogeneic therapy ^40–42^. NK cells also naturally mediate the killing of virally infected cells ^43–51^, including HIV-infected cells ^52–54^, both through receptor-mediated cytotoxicity and antibody-dependent cell-mediated cytotoxicity (ADCC) ^55,56^. Moreover, NK cells have been evaluated for their use as an HIV treatment ^52,57–59^, including in human clinical studies in which a single infusion of haploidentical NK cells plus either IL-2 or N-803 (an IL-15 superagonist^60^) was given to ART-suppressed people living with HIV (NCT03346499 and NCT03899480) ^57^. The NK therapy was well-tolerated and led to a decrease in the frequency of vRNA+ cells in lymph nodes. This effect may be further enhanced by engineering NK cells to express HIV-specific CAR molecules to direct their targeting and enhance activation ^32,61–63^, as several CAR NK clinical trials have been conducted with promising results in cancers ^64–66^, and preliminarily for HIV ^67–69^. In African Green Monkeys, which naturally control SIV and remain asymptomatic, innate NK cells have been shown to accumulate in lymphoid follicles at high levels and are naturally CXCR5+ ^19,70^. This finding supports the addition of CXCR5 to CAR NK cells for increased follicle-targeting capabilities.

There are a lack of animal models that recapitulate all components of HIV-1 infections in humans. SIV or SHIV-infected non-human primates are commonly used; however, their genetic diversity, particularly at immunogenetic loci, can determine their susceptibility to and maintenance of sustained infection ^2,3,71–76^. More recently, humanized immune system (HIS) mouse models have been used to study HIV pathology, vaccines, and therapeutics ^77^. DRAGA mice develop functional human B and T cells and a limited number of NK cells, sustain HIV infections, demonstrate B cell immunoglobulin class-switching, and elicit specific human cellular and antibody responses after vaccination ^78–85^. Unlike other mouse models, humanized DRAGA (hDRAGA) mice develop secondary lymphoid tissues with follicle-like structures (FLS) in lymph nodes and spleen ^84,86^. Importantly, HIV virions were shown to replicate in follicular-like CD20^high^ areas of secondary lymphoid tissues of HIV-infected hDRAGA mice ^84^. We have recently demonstrated that HIV-infected hDRAGA mice are a valuable model for studying immune cell therapies ^86^.

Given these findings, we engineered HIV-targeting CAR NK cells using non-viral DNA transposon technology and tested their safety and efficacy in infected hDRAGA mice. To further enhance the function of our CAR NK cell therapy, we armored our cells with soluble IL-15 and knocked out PD-1. Previous reports using innate immune effectors have demonstrated that cytokine armoring can enhance *in vivo* persistence, function, and resilience ^87,57,88^. Additionally, our group and others have demonstrated that knockout (KO) of PD-1 may increase the cytokine production and cytotoxic function of NK cells, as well as improve their persistence ^61,66,87,89^. CAR/CXCR5/IL-15/PD-1^KO^ NK cells (for brevity, hereon referred to as CAR NK cells) were evaluated for functionality compared to feeder-cell-activated control NK cells *in vitro* and *in vivo*. We found that a subset of CAR and control NK-treated animals had undetectable viral loads at study end. CAR NK-treated animals had higher total NK cell levels, with NK cells persisting up to 2 weeks longer than control NK-treated animals. Importantly, successful treatment was associated with faster viral rebound post-analytical treatment interruption (ATI), indicating the importance of treatment timing for NK cell therapies.

## Results

### CAR and control NK cells demonstrate killing efficacy against HIV-Env expressing cells *in vitro*

CAR NK cells were engineered using the *TcBuster* non-viral DNA transposon system to express a polycistronic construct encoding the CD4-MBL-CAR, CXCR5, and IL-15 with simultaneous PD-1 knockout using an adenine base editor (ABE) **(Fig. 1a).** As a control, unedited NK cells were cultured in the same conditions but were not engineered. Since there is variability between the functionality and expansion rates of NK cells from different donors^90^, NK cells from three deidentified human donors (numbered here as donors 10, 15, and 21) were engineered and evaluated in functional assays (**Fig. S1)**. Donor 21 was determined to be unsuitable for production due to poor viability of expanded control NK cells **(Fig. S1d)**, and as such, was not included in some assays. Despite this viability problem, subsets of CAR NK cells from all donors expressed CAR, CXCR5, and IL-15 at levels above the control cells **(Fig. S1a-b),** and donors 15 and 21 showed 96% and 99% knockout of PD-1, respectively **(Fig. S1c)**. In the presence of HIV-Env expressing cells, donor 10 CAR NK cells showed the highest levels of TNF-α, IFN-ɣ, and CD107a production **(Fig. S1e),** and donors 10 and 21 both showed slightly increased migration to CXCL13 compared to control NK cells **(Fig. S1f).** For these reasons, donor 10 was determined to be the most amenable to engineering and was selected for large-scale preparation of NK cells for infusion. A final round of *in vitro* functionality testing was performed on this infusion product **(Fig. 1a)**. The CAR NK cell product had 45% dual expression of CAR and CXCR5 **(Fig. 1b)** and a 74% (±23.7%) reduction in PD-1 expression compared to control NK cells **(Fig. 1c)**. Cell culture supernatant of CAR NK cells also contained 61.14 (±10.97) pg/mL of IL-15, versus 0 pg/mL from control NK supernatant **(Fig. 1d)**. CAR- and control NK cells were frozen after the second expansion. Both groups recovered from cryopreservation with moderate cell loss and expanded 27.3-fold or 22.7-fold, respectively, in the third and final expansion before use *in vivo* **(Fig. 1e)**. Finally, CAR and control NK cells both killed HIV-Env expressing target cells at high levels, with CAR NK cells having 80.7% (±10.4) specific lysis and control having 69.5% (±10.1) **(Fig. 1f)**. Taken together, these data indicate that CAR and control NK cells are efficacious *in vitro*.

**Figure 1:**
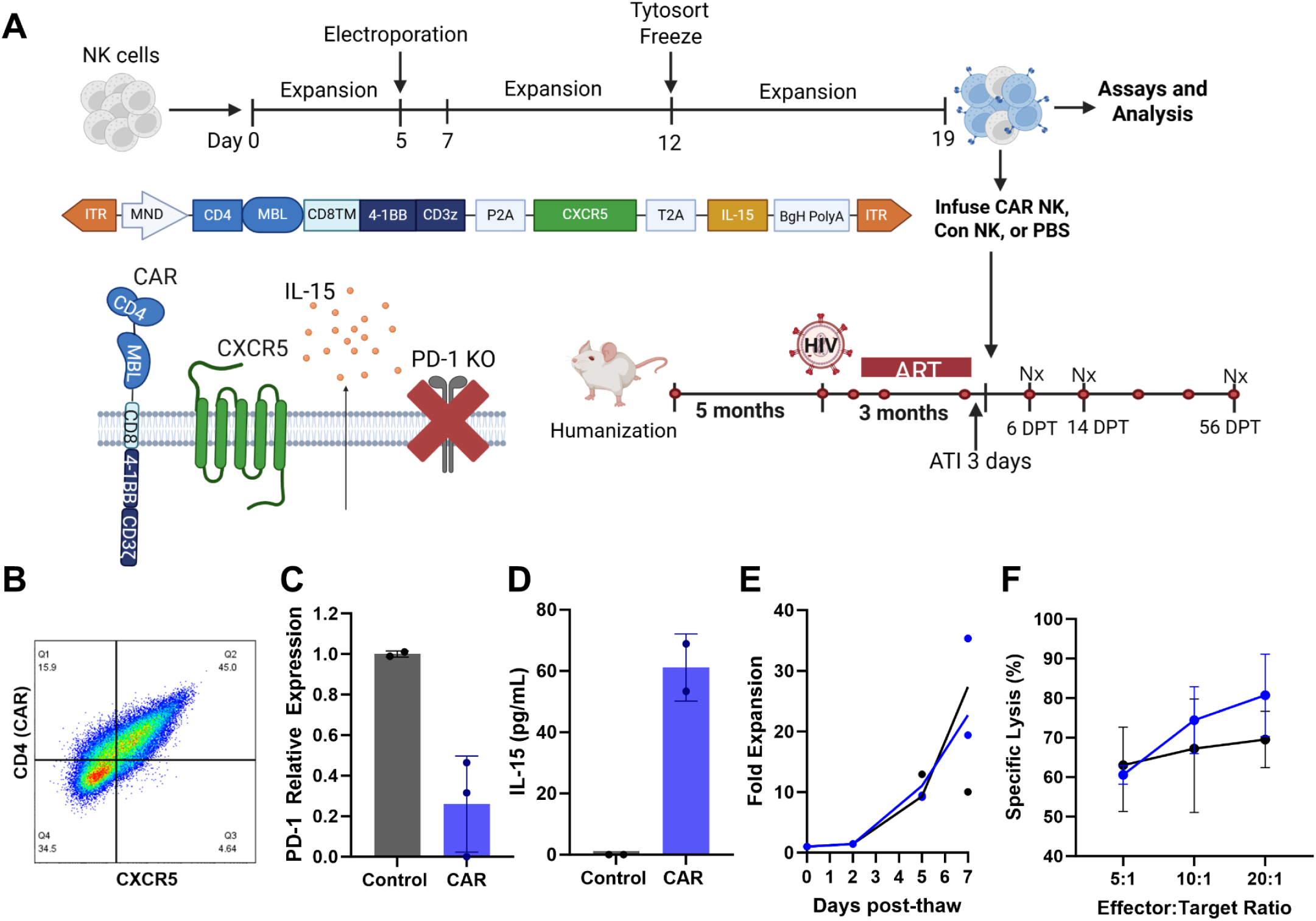
Study overview and *in vitro* testing of NK infusion products. A) Schematic figure showing the overview of NK cell production and the overview of the mouse study timeline. Red circles on the timeline indicate blood collection, black dots indicate scheduled necropsies. Schematic of transgene design: MND promoter, CAR (CD4-MBL-CD8TM-4-1BB-CD3z), CXCR5, IL-15, the BgH PolyA tail, with separation mediated by P2A and T2A sites, and inverted terminal repeats (ITRs). Schematic showing expression of all transgene components as well as base editor-mediated knock out of PD-1 on the NK cell surface. Figure made using Biorender.com. B) Expression of CAR (MBL) and CXCR5 molecules was confirmed in the cell product using flow cytometry. Cells were pre-gated sequentially on lymphocytes, singlets, live cells, CD56+, and CD3-. C) PD-1 knockout was detected at the RNA level by RT-PCR, CAR NK (blue) relative to control NK cells (black). D) IL-15 expression in CAR NK (blue) and control NK (black) cells via IL-15 ELISA of cell culture supernatant. E) CAR (blue) and control (black) cell proliferation over one week post-thaw. F) The specific lysis of HIV Envelope P815s by CAR (blue) or control (black) NK cells via DELFIA cytotoxicity assay. 5:1, 10:1, and 20:1 E:T ratios were tested. All assays were performed in duplicate or triplicate. Data are represented as mean + standard deviation (error bars).

### Treatment with CAR or control NK cells at the time of viral recrudescence led to control of HIV viremia

After confirming CAR and control NK cells function *in vitro,* the safety and efficacy of these therapies were evaluated *in vivo* with HIV-infected hDRAGA mice. Animals were sorted into infusion groups taking into account their age, weight, time post-humanization, peak viral load, viral load area under the curve, and CD4 counts pre-infection **(Fig. S2)**. The DRAGA mice were infused intravenously with CAR NK cells or control NK cells at a dose of 1×10^5^-1.5×10^5^ cells/g or with saline buffer (PBS) solution. Viral loads were monitored throughout the study as the primary marker of therapeutic efficacy. HIV viral peak was observed in all mice by 2 weeks post-infection **(Fig. S3a-c)**. Mice were then given ART-containing chow from which they could free feed. Viral control was obtained in all mice between 4 and 8 weeks post ART onset **(Fig. S3a-c)**. It was initially observed that mice were not eating the ART chow readily when it was placed in the hopper and this led to a delay in viral control; however moving the ART chow to paper cups on the base of the cage led to increased uptake and viral control in all animals. Once all mice were below the detection limit for HIV (less than 200 HIV RNA copies/mL of peripheral blood), they were infused with CAR NK, control NK, or PBS at 3 days post-ATI. In all animals, viral loads increased back to pre-ART levels between 3 and 28 days post-removal from ART **(Fig. 2a-g, Fig. S3a-c)**. At 28 days post-treatment (DPT), 4 of 8 CAR NK-treated animals and 2 of 4 control NK-treated animals had undetectable viral loads, compared to 0 of 6 in the PBS control group (Fisher’s Exact test, P=0.1070) **(Fig. 2g).** Between 42 and 56 DPT, one additional animal in the CAR NK treated group declined to undetectable viral loads **(Fig 2i)**. At study end, 62.5% (5/8) of CAR-treated, 50% (2/4) of control-treated, and 0% (0/6) PBS-treated animals had undetectable viral loads **(Fig. 2h-i).** The differences between the number of animals that controlled between the CAR NK, control NK, and PBS group did not quite meet the criteria for statistical significance with the Fisher’s exact test (P=0.0546). Importantly, however, there were undetectable viral loads in 58% (7/12) of NK-treated animals (CAR and control) compared to 0% (0/6) in the PBS group demonstrating that overall, NK cell treatment led to significant reductions in viral loads (P=0.0377) **(Fig. 2j).** Interestingly, in the animals that had undetectable viral loads by 56 DPT (controllers), the time to viral rebound following ART interruption was on average 15.5 days shorter compared to non-controllers (P=0.0278) **(Fig. 2k)**. On average, it took only 3.7 days for controllers to show detectable viremia following ATI, while it took 19.2 days for non-controllers. Thus, in all animals that controlled infection, viral rebound was earlier and closer to the time of therapeutic cell infusion. These data indicate that the NK cell treatments were only effective when the timing of cell infusion aligned with viral recrudescence.

**Figure 2:**
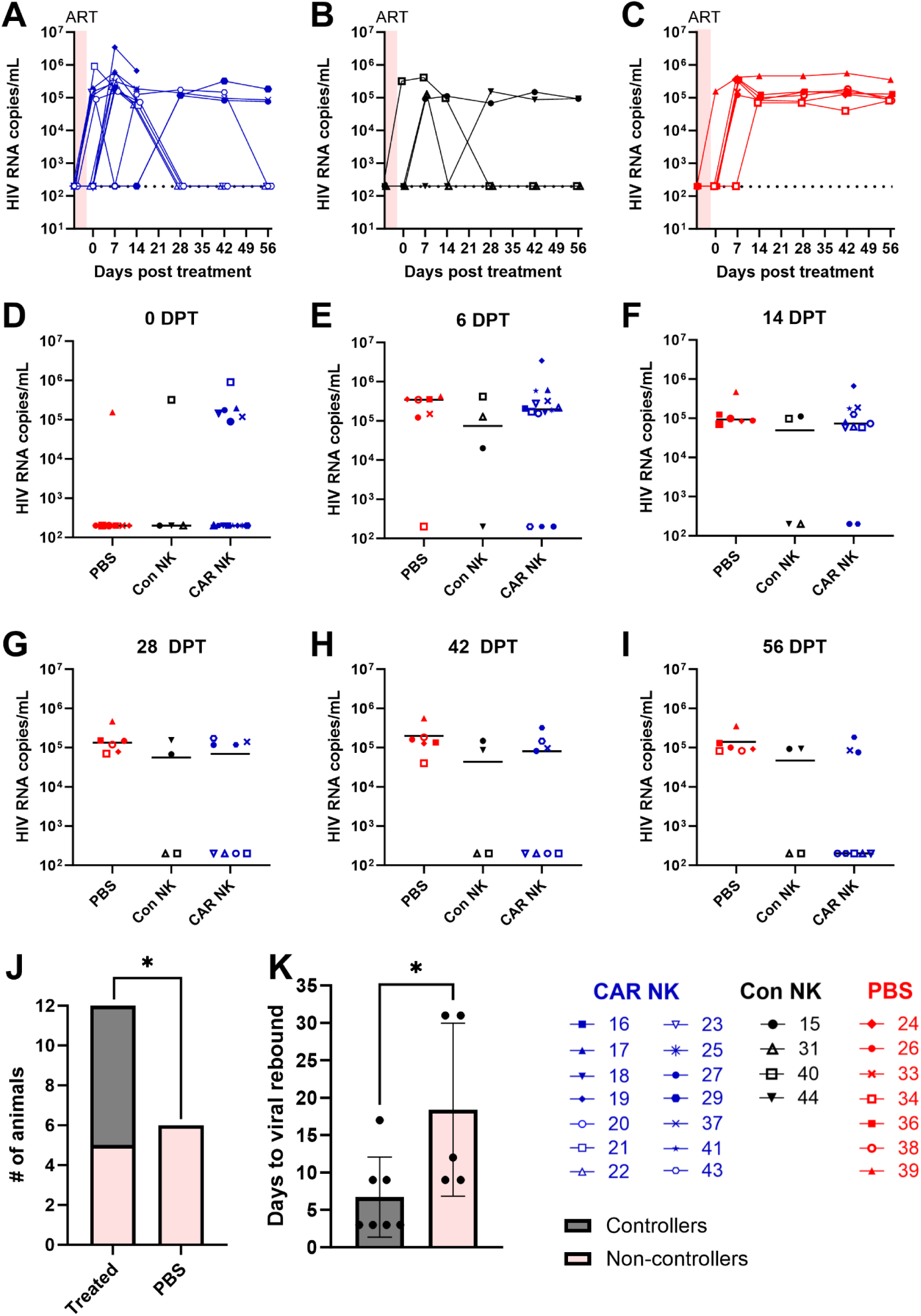
Post-treatment viral loads. Viral loads over time in A) CAR NK treated animals (blue), B) control NK treated animals (black), and C) Viral loads in CAR (blue), control (Black), and PBS (red) treated groups at D) 0 DPT, E) 6 DPT, F) 14 DPT, G) 28 DPT, H) 42 DPT, and I) 56 DPT. Lines represent median values. J) Number of animals in either the treated (CAR and control NK) or untreated (PBS) groups at 56 DPT, with the percentage of controllers (gray) or non-controllers (pink) at 56 DPT (P=0.0377, Fisher’s exact test of proportion within treated versus PBS groups). K) Controllers (gray) or non-controllers (pink) at 56 DPT by number of days to viral rebound post-ART interruption (P=0.0278, Mann-Whitney test). Mean values with SD are shown.

### CAR and control NK cell treatments are safe and not associated with any adverse health outcomes *in vivo*

Animal health and well-being were monitored for the duration of the study by visual inspection and weight. Of the original 30 animals, six acquired severe adverse outcomes (**Table S1)** and were euthanized prior to the intended study endpoint. Five of these animals were euthanized prior to being assigned treatment groups, and one additional mouse was euthanized after assignment but was in the PBS-treated group (**Table S1**). In all cases, observed symptoms were qualitatively consistent with chronic graft-versus-host disease (GVHD) in humanized mice ^91^ **(Table S1**). We can therefore conclude that there were no adverse symptoms associated with receiving CAR or control NK cells in this study.

### CAR NK treatment may temporarily increase CD4:CD8 T cell ratios

CD4+ T cell levels were monitored as a secondary measure of treatment outcome, as decreasing CD4 T cell levels are associated with HIV progression, and CD4 maintenance is associated with therapeutic efficacy^92^. Here, we opted to use CD4:CD8 ratios as opposed to CD4 T cell counts, as ratios were not impacted by the fluctuations in overall lymphocyte numbers which were observed in this study (**Fig. 4i**). One week after HIV infection, 36% of infected hDRAGA showed a decline in CD4:CD8 ratios from baseline, and this percentage increased to 87.5% by 4 weeks post-infection **(Fig, S3d-f)**. During ART, 67% of animals showed maintenance or an increase of CD4:CD8 ratio, with the remaining animals showing a reduction in the rate of CD4:CD8 decline **(Fig. S3d-f)**. Following infusion, 75% of CAR NK-treated, 33% of control NK-treated, and 43% of PBS-treated animals showed an increase in CD4:CD8 ratios at 6 DPT (P=0.1790) **(Fig. 3a-e).** The magnitude of the increase also differed between groups, with CAR NK-treated animals showing a median 1.74-fold increase, compared to 0.96-fold in control-treated and 0.88-fold in the PBS group. We were unable to obtain data for DRAGA 15 at the pre-treatment time point, so this animal was excluded from these analyses. Despite the initial increase, the effect was diminished in all animals by 14 DPT, and CD4:CD8 ratios remained at low levels for the remainder of the study **(Fig. 3e-i).** These data indicate that CAR NK treatment may have led to a temporary increase in CD4 T cells above control levels following treatment.

**Figure 3:**
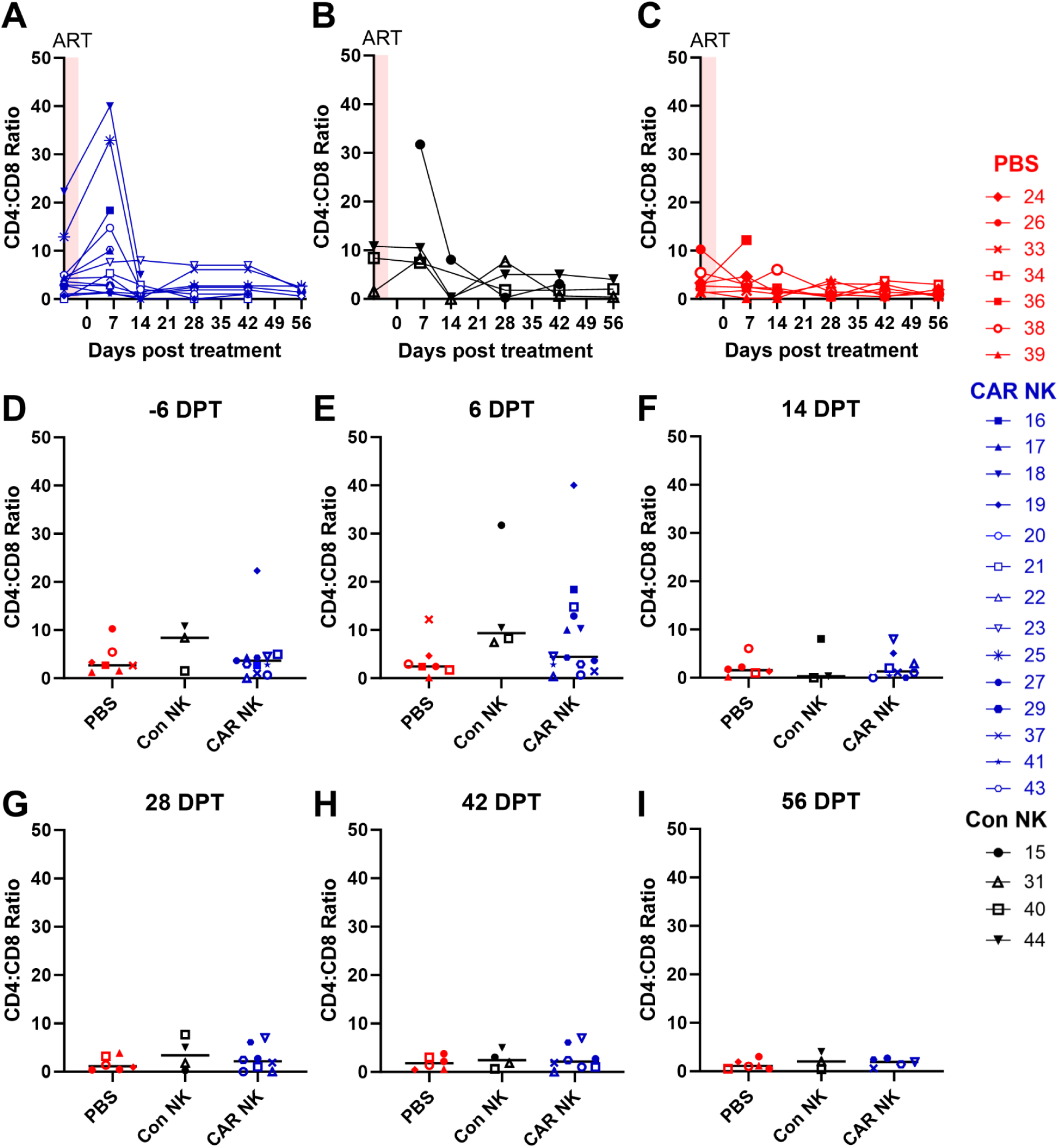
Post-treatment CD4:CD8 ratios. CD4:CD8 ratios over time in A) CAR NK-treated animals (blue), B) control NK-treated animals (black), and C) PBS-treated animals (red). CD4:CD8 ratios in CAR (blue), control NK (black), and PBS (red) treated groups at D) -6 DPT, E) 6 DPT, F) 14 DPT, G) 28 DPT, H) 42 DPT, and I) 56 DPT. Lines represent median values. CD4:CD8 ratios were determined by flow cytometry and pre-gated on Lymphocytes, Singlets, Live Cells, Human CD45+, Mouse CD45-, CD3+ (see gating strategy in **Fig. S4**).

**Figure 4:**
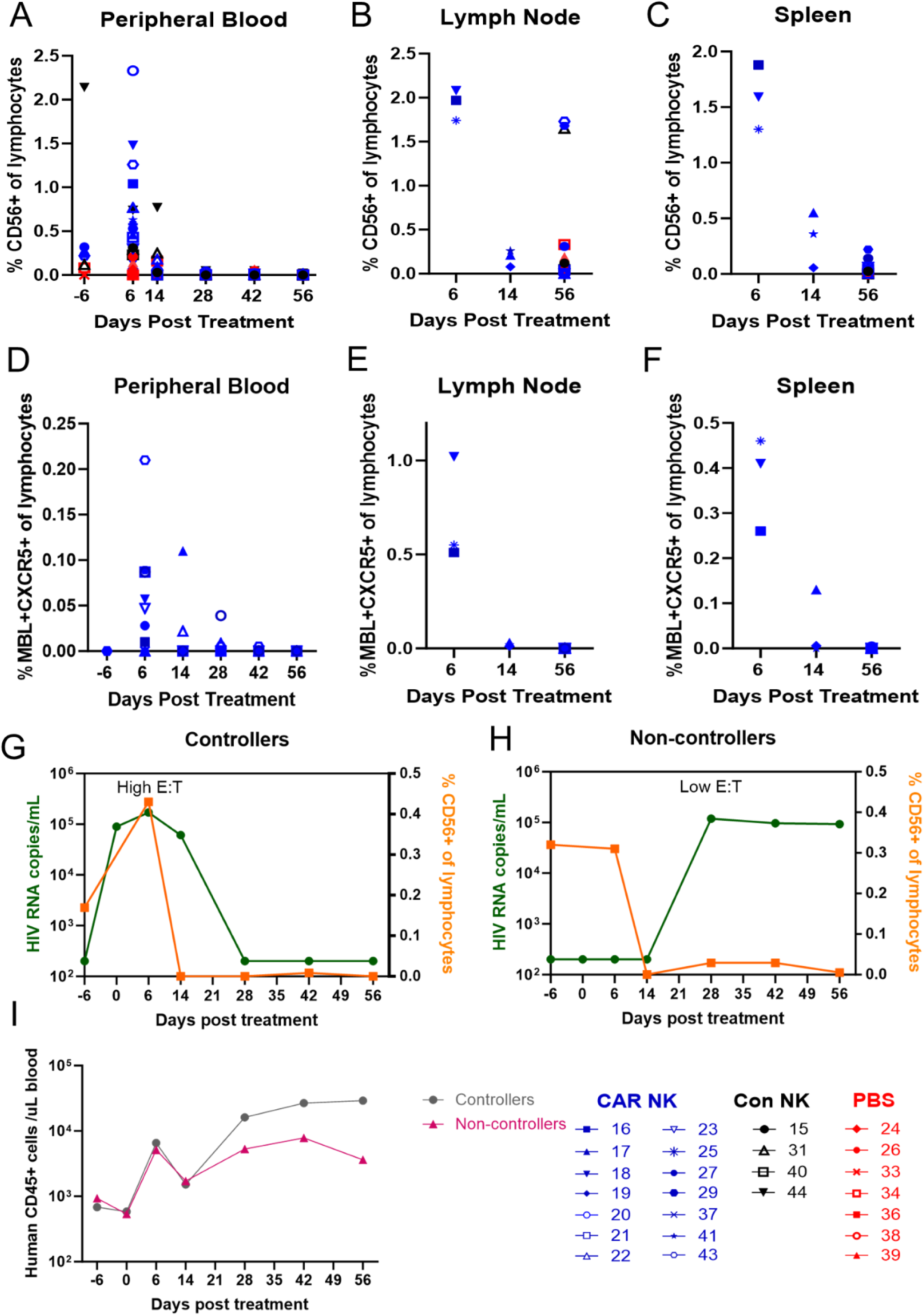
CAR NK levels post-treatment. NK (CD56+) cells in CAR (blue), control (black), or PBS (red) treated groups from A) Peripheral Blood, B) Lymph node (at necropsies), and C) Spleen (at necropsies). CAR+CXCR5+ NK cells over time in D) Peripheral Blood, E) Lymph node (at necropsies), and F) Spleen (at necropsies). Flow was pre-gated on Lymphocytes, Singlets, Live Cells, human CD45+, mouse CD45-, CD3+, and then through CD56+ MBL+ CXCR5+ (see **Fig. S4** for gating strategy). Median viral loads (green), NK cell levels (orange), post-treatment in G) controllers or H) non-controllers. I) Median human hematopoietic cell counts (CD45+)/ ul blood of controllers (gray) or non-controllers (pink).

### Peak NK cell concentration at the time of viral recrudescence is associated with viral control

Following infusion, DRAGA mouse peripheral blood samples were regularly assessed for persistence of NK and CAR NK cells, and spleen and lymph node samples were assessed at each necropsy time point. In the peripheral blood, NK cell levels peaked at 6 DPT for both the CAR and control NK treated groups, with the CAR NK-treated animals having 1.88 times higher concentration of NK cells than control NK-treated animals on average **(Fig. 4a).** NK cells in CAR NK treated animals were detected at low levels until 28 DPT, while control NK cells were only detected until 14 DPT **(Fig. 4a)**. In the spleen and lymph nodes of CAR and control NK treated animals, NK cell levels were slightly lower but similarly peaked at 6 DPT **(Fig. 4b).** In all but three animals, NK cells declined to background levels prior to 56 DPT **(Fig. 4b and c).** CAR/CXCR5+ NK cells were detected at low levels in the peripheral blood, with peak concentration occurring at 6 DPT **(Fig. 4d)**. Low levels of CAR/CXCR5+ NK cells were detected in the spleen and lymph node at 6 DPT and declined to undetectable levels by 14 DPT in all but one animal’s lymph nodes **(Fig. 4 e and f).**

In animals which controlled viral loads, median peak NK cell concentration timing coincided with median peak viral loads post-ATI, resulting in relatively high *in vivo* effector NK cell to target cell ratios (E:T) at viral peak **(Fig. 4g)**. Conversely, in noncontrollers, median NK cell concentration diminished before viral rebound resulting in relatively low *in vivo* E:T at viral peak **(Fig. 4h).** Controllers did have a 1.39 times higher median peak NK cell levels compared to non-controllers; however, the observed difference was not statistically significant (p=0.3649) (**Fig. 4g-h).** Taken together, these data indicate that NK cell levels at the time of viral rebound may dictate viral control.

To determine a potential mechanism for the sustained viral control even as NK cell levels decline, we analyzed the human hematopoietic (CD45+) cell response after treatment in the responder and non-responder groups **(Fig. 4i)**. All animals had increasing levels of human hematopoietic cells at 28 DPT independent of the influx caused by treatment with NK cells, with this increase being somewhat more pronounced in the controller group. The hematopoietic cell levels remained high for the duration of the study in the controller group, while levels began to decrease in the non-controllers. At 56 DPT, human hematopoietic cells of the controllers were 8.07-fold higher than those of the non-controllers, indicating that the humanized immune response in the DRAGA mice may have been sufficient to take over and maintain control after effective intervention with NK cell therapies.

Spleen tissue sections from the three animals at 6 DPT were stained using RNAscope and immunohistochemistry to detect CAR NK cells, HIV vRNA, and CD20 (B cell marker used to delineate FLS). The three animals were found to have variations in their humanization and, therefore, in size and abundance of their FLS. Animals with high levels of humanization showed more FLS than animals with low levels of humanization. In sections with FLS, the HIV vRNA+ cells were concentrated within the FLS **(Fig. S5).** In the sections examined from 6 DPT spleens, no CAR+ cells were detected above background levels (data not shown).

## Discussion

Here, we demonstrate additional evidence of the promise of NK therapies for treating HIV infection. Our findings indicate the importance of timing the delivery of therapeutic cells with the timing of viral rebound. We found that both CAR NK cell and control NK cell treatments showed efficacy at suppressing HIV infections when infused at the time of viral recrudescence following ART interruption. In this study, 63% (5/8) of CAR NK-treated and 50% (2/4) control NK-treated animals had undetectable viral loads by 56 DPT compared to 0% (0/7) saline-treated animals. There was a statistically significant difference in the number of animals that controlled HIV between the NK treatment groups (CAR and control NK) and animals that received saline treatment (P=0.0377). Interestingly, we also observed a statistically significant difference in the time to viral rebound between animals that controlled and did not control infection (P=0.0278). In animals that controlled infection, median viral rebound occurred at the time the therapeutic cells were infused, resulting in relatively high *in vivo* E:T ratios at viral peak. Conversely, in animals that did not control infection, in most cases viral rebound occurred after the infused NK cells declined to very low levels, resulting in relatively low *in vivo* E:T at viral peak. These findings indicate that NK treatment at the time of viral recrudescence is important for the success of the treatment.

Despite NK cell levels diminishing by 14-28 DPT, animals that controlled HIV sustained undetectable viral loads until 56 DPT. This finding suggests that NK cells provided initial management of viral loads and allowed time for the endogenous adaptive immune response to rebound. This theory is supported by the finding that human CD45+ cells proliferated in all treatment groups at 28 DPT. Once rebounded, the adaptive immune response may be better equipped to maintain control of HIV replication in animals with successful NK intervention, as controllers maintained higher hematopoietic cell levels for the duration of the study, while non-controller levels began to decrease. This aligns with our previous studies where we found sustained control of SIV in CAR/CXCR5 T cell-treated animals, long after the CAR T cells became undetectable^31^. Thus, long-term control of HIV may be achieved with relatively short-acting cellular therapies.

Additional doses of CAR NK or control NK cells could lead to improved viral suppression in the animals that did not control HIV. Since timing NK cell treatment with viral rebound seems to be important for viral control, infusing a second dose of NK cells could help improve the odds of treating at the time of viral rebound in all animals, including those with delayed viral rebounds. This theory aligns with NK cell therapies to treat myeloma, where multi-dosing regimens were shown to improve NK treatment efficacy^93–95^. In addition to multiple doses of NK cells, an earlier dose may lead to viral suppression without viral rebound occurring. Indeed, Kim et al. found that treating HIV infected mice with allogeneic peripheral blood-derived NK cells one and six days after ART interruption led to a significant delay in viral rebound and significantly lower numbers of rebound barcode HIV variants ^58^.

We found no evidence of adverse outcomes related to NK cell treatment. Like most humanized mouse models, DRAGA mice risk developing chronic GVHD ^96^. It is of interest that the 6 animals that had GVHD-like symptoms were all female and were humanized with male donor cells. Although it is widely considered safe to transplant female NIS mice with male stem cells ^97,98^, some studies in human stem cell transplant have indicated that sex-mismatched stem cell engraftments can lead to increased risk of developing GVHD ^99,100^. Additionally, 5 of the 6 mice were humanized with cells from the same human stem cell donor, and donor-specific factors have been known to increase the risk of developing GVHD ^101^. Nevertheless, all adverse symptoms occurred in either pre-treatment or PBS-treated groups in this study, and we found no evidence that either the CAR or control NK therapies caused any adverse outcomes.

In some animals, especially in the CAR NK group, cell infusion led to a temporary increase in CD4:CD8 ratios, indicating a rise in CD4 T cell levels following treatment. The magnitude of this difference was greatest in the CAR-treated group; however, this increase was not statistically significant. It is possible that this increase in CD4:CD8 ratio was caused by antigen resurgence after release from ART, as this could have led to a temporary increase in immune activation and cell expansion, before resuming CD4 loss ^102^. It is also possible that the increased cytokine production by the CAR NK cells, including soluble IL-15, led to a further increase in immune activity when compared to control NK or PBS-treated groups.

We detected low levels of CAR NK cells in disaggregated spleen and lymph nodes from treated animals by flow cytometry, with the highest levels detectable at 6 DPT. However, no CAR+ cells were detected by RNAscope in spleen sections that were examined. Taken together, these data indicate that while the therapies showed efficacy in this model system, they did so without accumulating to high levels in secondary lymphoid organs.

NK cell subsets express CXCR1, CXC3R1, and ChemR23, and, at low levels, CCR7, allowing them to migrate to lymphatic tissues in inflammatory conditions ^103^, like HIV infection ^104,105^. To further increase the levels of NK cells migrating to these lymphatic tissues, in future studies, we might increase the expression of CCR7 and L-selectin on PB-NK cells, which may lead to increased migration to lymphatic tissues. Transient expression of CCR7 combined with CXCR5 may be ideal for B-cell follicle homing, as other cell types like T Follicular Helper cells use CCR7 to enter lymphatic tissues and then downregulate CCR7 and rely on CXCR5 to enter follicles ^106–108^. Somanchi et al. have previously shown that NK cells can be made to transiently express CCR7 via trogocytosis following co-culture with CCR7-overexpressing K562 feeder cells ^109^. Using this method, CCR7 expression returned to baseline levels after 72 hours. Alternatively, mRNA transfection could be used to express CCR7, L-selectin, and CXCR5, which have been shown to improve homing of NK cells to the lymph nodes ^110^. Future studies with NK cells engineered with CCR7 and CXCR5 are warranted to determine whether these homing molecules are sufficient to clear the HIV reservoirs in lymphatic tissues.

Additional strategies to achieve better long-term control of infection include CAR NK or NK cell immunotherapy, combined with other strategies aimed at eliminating the HIV/SIV reservoir. Such therapies might include latency reversal agents to reactivate latent HIV infected cells ^58,111–113^, BIKES or TRIKES to connect immune cells with HIV expressing cells^114–116^, or agents that modify the immune system, such as an IL-15 superagonist ^57,117^ or rapamycin^118,119^.

Taken together, these studies demonstrate the promise of NK cell therapies for treating HIV. We showed that CAR and control NK cells are safe and lead to undetectable viral loads in some treated animals. Importantly, we uncovered a potential mechanism of therapeutic effect, suggesting that the timing of treatment with viral recrudescence was key to efficacy. Future studies could explore multiple rounds of NK cell infusions to achieve superior efficacy. With refinement of treatment timing, immunotherapeutic NK cells could lead to long-term suppression of HIV without the use of life-long ART.

## Materials and Methods

### Production of engineering reagents

ABE8e plasmid was obtained from Addgene (https://www.addgene.org/138489/) and cloned into a pmRNA vector. ABE8e mRNA was produced by Trilink Biotechnologies. Nanoplasmid encoding the bispecific CD4-MBL CAR with the 41BB intracellular domain, CXCR5, and IL-15 was synthesized by Aldevron. Specific sequence included in **Fig. S6.**

### Donor NK Cell Isolation and Expansion

Peripheral blood mononuclear cells (PBMCs) from three de-identified healthy human donors (labeled as donors 10, 15, and 21 for the purpose of distinguishing) were obtained via automated leukapheresis (Memorial Blood Centers, Minneapolis, MN, USA) and further isolated using a Ficoll-Hypaque (Lonza) gradient. NK (CD56+/CD3-) cells were isolated using CliniMACS, following the manufacturer’s instructions. NK cells were cultured in CTS AIM V SFM (ThermoFisher) supplemented with 5% CTS Immune Cell SR (Gibco), Penicillin/Streptomycin, and IL-2 (100 IU/mL), as previously described ^61^. Activation of NK cells was achieved by co-culturing NK cells with X-irradiated (100 Gy) feeder cells (K562 cells expressing membrane-bound IL-21 and 4-1BBL) at a 1:2 ratio of NK cells to feeders. Cells were supplemented with additional medium and IL-2 up to 100 IU/mL every 2-3 days.

### Production of CAR/CXCR5/PD-1KO/IL-15 NK cells

CAR NK cells were engineered using the *TcBuster* non-viral DNA transposon system to express a polycistronic construct encoding CAR, CXCR5, and IL-15. An adenine base editor and a guide RNA complementary to the PD-1 gene (CACCTACCTAAGAACCATCC) were utilized to knock out PD-1 expression. Isolated NK cells were expanded for 5 days following the culture and activation protocol above. Cells were electroporated using the MaxCyte (ThermoFisher) with transposase mRNA, nanoplasmid DNA, ABE8E mRNA, and guide RNA as previously described ^64^, or electroporated with no additional reagents for control NK cells. After a 2-day recovery, CAR NK cells were sorted on CD56+ CD4+ using the MACSQuant Tyto Cell Sorter (Miltenyi Biotec) and expanded for an additional 7 days following the procedure described above.

### Cell culture of target cells

Inducible truncated HIV-Env P815s-The P815 truncated HIV Env cell line was produced by retronectin-mediated transduction of P815 cells (a gift from Dr. Geoffery Hart) using a lentivirus produced from a tet-inducible truncated HIV Envelope plasmid (kindly provided by Dr. Alon Herschhorn; Tet-One Inducible Expression System, Clontech) and psPax2 and VSV-G plasmids. Puromycin-resistant cells were cloned by limiting dilution to produce a stable cell line. Cells were cultured in DMEM + 10% tet-free FBS + 1% P/S and maintained at a density of 0.7e5-1e6 cells/ml medium. Selective pressure for cells containing the inducible HIV-Env construct was maintained by adding 1 μg/mL puromycin to the culture. HIV Env expression was induced by adding 1 μg/mL of doxycycline to the culture for 72 hours. Expression of HIV Envelope was assessed by flow cytometry.

Trunc-HIV Env expressing PBMCs-The HIV Envelope gene, truncated 145 amino acids at the C terminus, was inserted into the pMSGV gammaretroviral plasmid by Gene Universal. Pseudotyped gammaretrovirus containing truncated HIV-Env was produced by Lipofectamine-mediated transfection of 293T cells cotransfected with pBS-CMV-gagpol, RD114, and pMD.g, as described previously ^29^. PBMCs from a human donor were transduced with HIV-Env pseudotyped gammaretrovirus as previously described ^29,120^. As an assay control, NK cells were subjected to the same culture conditions with no virus present. HIV-Env PBMCs were evaluated for expression and frozen for future expansion and use. Prior to running assays, cells were thawed and expanded in a GREX with 50 IU IL-2 + 55 μM beta-mercaptoethanol (Gibco) added to the medium for an additional 4 days. Expression of HIV Envelope was assessed on a Cytoflex flow cytometer after gating on alive, single cells.

### Pro-inflammatory cytokine and degranulation marker assay

Antigen-specific production of TNFα, IFNγ, and CD107a were assessed by coculturing CAR NK cells (effector) with either Trunc-HIV-Env PBMCs or Trunc-HIV-Env P815s at an E:T ratio of 1:1. Anti-human CD107a (Biolegend, H4A3, BV421) was added, and cells were incubated for 1 hour at 37 °C, at which point brefeldin A (Biolegend) and monensin (Biolegend) were added, and cells were incubated for 3 hours at 37 °C. Intracellular cytokine staining was performed for TNF-α (Biolegend, MAb11, BV650) and IFNγ (Biolegend, B27, PerCP/Cyanine 5.5) prior to surface staining for MBL (Invitrogen, 3E7), CXCR5 (Invitrogen, MU5UBEE, PE), and CD56 (Immunotech, N901, PC5.5). Samples were acquired on a CytoFLEX (Beckman) flow cytometer and analyzed via FlowJo v 10.2 software (Becton Dickenson). Cells were gated on lymphocytes, single cells, CD56+ MBL+ CXCR5+, and then TNF-α+, IFN-γ+, or CD107a+.

### Cell cytotoxicity assays

The DELFIA EuTDA assay (Revvity) was used to assess cell cytotoxicity per the manufacturer’s instructions. Briefly, HIV-Env-expressing P815 cells were labeled with BATDA for 30 minutes at 37 °C and subsequently washed with PBS. CAR NK cells or control NK cells were plated with HIV-Env or WT P815s at 5:1, 10:1, and 20:1 E:T ratios. After 4 hours of incubation, 20 μL of supernatant was added to 200 μL of Europium solution and transferred to a flat-bottom plate, and time-resolved fluorescence was measured using a Synergy 2 (Biotek). Spontaneous and maximum release wells were prepared by adding BATDA-loaded P815s to a well containing medium alone or medium with 10% lysis buffer, respectively. Specific release was defined as (Experimental release-spontaneous release)/(maximum release-spontaneous release).

### Analysis of PD-1 expression

Expanded CAR and control NK cells were harvested, and total RNA was isolated using the RNeasy Mini Kit (Qiagen, 74104) following the manufacturer’s instructions. 1 µg of RNA was reverse-transcribed into cDNA with the SuperScript IV RT Kit (Life Technologies, 18090050). qPCR reactions were performed using SsoAdvanced Universal SYBR® Green Supermix (Bio-Rad, 1725274) with the following primers: PD-1 forward (CCCTGGTGGTTGGTGTCGT) and reverse (GGCTCCTATTGTCCCTCGTGC); β-actin forward (AGGCACCAGGGCGTGAT) and reverse (TGGGGTACTTCAGGGTGAGGA). Amplification was carried out on a CFX96 Real-Time PCR System (Bio-Rad), and data were analyzed using CFX Manager 3.1 software (Bio-Rad, CA, USA). Relative gene expression was calculated using the comparative Ct (ΔΔCt) method with β-actin as the internal control, and the percentage reduction in expression was calculated as: %Reduction=(1−Relative Expression)×100.

### Detection of IL-15 production

CAR and control NK cells were collected following their third expansion. Cells were centrifuged, and the supernatant was collected. Human IL-15 ELISA kits (Abcam) were used to determine IL-15 concentration in the supernatant following the manufacturer’s instructions.

### Migration assay

Migration of CAR NK cells towards specific human cytokines was conducted as previously described ^29,120,121^. Briefly, one million CAR or control NK cells (in 100 µl of X-VIVO 15 with 0.1% BSA) were placed in the upper chamber of a Transwell plate (Costar) with a 5.0-µm membrane. The lower chamber was filled with X-VIVO with 0.1% BSA alone or X-VIVO with 0.1% BSA and the chemokine human CXCL13 (Peprotech) (2.5 µg/ml). After a 4-hour incubation at 37°C, the cells were collected and counted using a CytoFLEX flow cytometer (Beckman Coulter). AccuCheck counting beads (Invitrogen) were added to each sample to ensure accurate cell counts. Specific migration was defined as (Cells that migrated to CXCL13-cells that migrated to medium alone)/input cells.

### HIV infection and treatment of DRAGA mice

DRAGA mice (HLA-A2. HLA-DR4. RAG1 KO. IL-2R g c KO. NOD) are on an NRG background and have transgenes for HLA-A2 and HLA-DR4, allowing for enhanced donor-specific engraftment with CD34+ human hematopoietic stem cells from cord blood ^78^. They were bred and humanized at the Veterinary Service Program at WRAIR/NMRC before being transported to the University of Minnesota. 30 humanized DRAGA (hDRAGA) mice (18 male and 12 female), humanized from two human donors, were housed under specific pathogen-free conditions. Following a one-week quarantine period, hDRAGA mice were evaluated for the reconstitution levels of human immune cells and then infected intraperitoneally with 20,000 tissue culture infectious dose 50 (TCID50) of HIV-1Ba-L in 200 µL PBS, as previously described ^122^. Four weeks after infection, the mice were switched from standard feed to 1/2″ pellets of irradiated Teklad chow 2020X containing 1500 mg emtricitabine, 1560 mg tenofovir disoproxil fumarate, and 600 mg raltegravir per kg (Research Diets, New Brunswick, NJ), as described by Tsai et. al. ^123^ in a hopper. It was initially observed that mice were not eating the ART chow readily when it was placed in the hopper, so the ART chow was instead moved to paper cups on the ground. Once all mice had undetectable HIV viral loads, they were switched back to standard Picolab dry feed. Animals were sorted into infusion groups taking into account their age, weight, time since infusion, peak viral load, viral load area under the curve, and CD4 counts pre-infection **(Fig. S2)**. Three days post-ART Interruption (ATI), the mice received an intravenous administration of CAR NK cells, control NK cells, or PBS at a dose of 1×10^5^-1.5×10^5^ cells/g. Animals were euthanized at 6, 14, and 56 days post treatment (DPT) via carbon dioxide inhalation followed by cervical dislocation. Mice were visually inspected daily and weighed weekly to monitor health and well-being.

### Blood and tissue collections from hDRAGA mice

Blood samples (up to 150 µL) were collected from the facial veins with the following frequencies: weekly to monitor viral infection, once, 8 weeks following ART initiation, after infusion on days 6, 14, and then biweekly until day 56. The blood samples were separately processed for isolating PBMCs and plasma, as previously described ^86^. Spleen and lymph nodes were collected from animals at the time of necropsy (6, 14, and 56 DPT) and divided into two parts. One part was disaggregated and lysed to remove red blood cells before being processed as described below for flow cytometric analysis. The remaining spleen and lymph node were fixed in 4% PFA and embedded in paraffin blocks by the U of M Clinical and Translational Science Institute and sectioned at 5 μm as previously described ^31^

### Whole blood cell counts and phenotype analysis

Cell counts were performed using 50 µL of whole blood, as described previously ^86^. Briefly, red blood cells were lysed using ACK lysing buffer (Gibco), and samples were washed. The cells were resuspended in 150 µL of PBS, and 50 µL of this suspension was mixed with a known concentration of AccuCheck counting beads (Invitrogen). The remaining resuspended cells were stained for flow cytometry as described below.

### Flow Cytometry of hDRAGA samples

Samples collected from spleen and lymph nodes were disaggregated and lysed. Samples from peripheral blood, spleen, and lymph nodes were then characterized using antibodies to stain for the following epitopes: CD4 (BD biosciences, M-T477, FITC), CXCR5 (Invitrogen, MU5UBEE, PE), MBL (Invitrogen, 3E7) conjugated to Alexa Fluor 647, CD3 (BD Biosciences, SP34.2, AF-700), CD56 (Immunotech, N901, PC5.5), and CD8 (Invitrogen, 3B5, PacBlue). Anti-mouse CD45 (Biolegend, 30-F11, BV785) and anti-human CD45RA (BD Biosciences, 5H9, PE-Cy7) were used to distinguish mouse and human hematopoietic cells. Samples were acquired on a Beckman Coulter CytoFlex cytometer, and a minimum of 10,000 events were collected for each sample. All data were analyzed using FlowJo software version 10.2 (Becton Dickinson).

### Plasma HIV-1 viral load determination

Plasma viral loads were detected as previously described ^86^. Up to 50 µL of plasma was used for RNA extraction using a QIAamp Viral RNA Mini Kit (QIAGEN) according to the manufacturer’s instructions. HIV-1 vRNA quantitation was carried out using real-time PCR with an HIV Quantitative TaqMan RT-PCR Detection Kit (Norgen Biotek Corp.) on a CFX96 Real-time PCR System (BioRad). Data collection and analysis were performed using CFX Manager 3.1 software (Bio-Rad, CA, USA), as previously described ^86^.

### RNAscope and immunohistochemistry

RNAscope *in situ* hybridization (ACD) and immunofluorescence were performed following manufacturer specifications and as previously described ^31^. Briefly, 5μm tissue sections were deparaffinized, pre-treated, washed, dehydrated in absolute ethanol, and air-dried. Sections were incubated with pre-warmed premixed target probes targeting HIV-Env and the CD4-MBL region of the CAR molecule and amplified. Opal 570 and Opal 650 were used for HIV-Env and CD4-MBL, respectively. For immunofluorescence staining, sections were washed, blocked, and incubated with primary rabbit anti-human CD20 antibody (Abcam, EP459Y) at 1.12 μg/mL. Sections were washed and incubated with 10 μg/mL Alexa 488-labeled goat anti-rabbit secondary antibody (Invitrogen) diluted in blocking buffer for 1 hour at RT. After washing, sections were counterstained with 1μg/mL DAPI and mounted in Prolong Gold (Invitrogen).

### Quantitative image analysis for RNAScope

Sections were imaged using a Nikon Ti-E confocal microscope. FLS and extrafollicular areas were delineated by morphology, with FLS identified as clusters of closely aggregated CD20+ cells. Qupath software was used to perform cell counts, delineate tissue areas, and define CAR+DAPI+ and HIV+DAPI+cells above background expression levels on negative-stained tissues. For these analyses, for each animal, 3 tissue sections were examined, and all FLS (if any existed) were examined in each tissue section.

### Statistical analysis

All statistical analyses were performed using GraphPad Prism version 10. Mann-Whitney or Fisher’s exact tests were used where appropriate. Standard deviations or ranges are indicated by the ± symbol and specified in the text where appropriate.

## Data availability statement

The data underlying this article will be shared on reasonable request to the corresponding author.

## Acknowledgments

This work was supported by amfAR Target (110411) grant to P.J.S., V.V., and B.S.M.; NIAID R01 (AI161017) to P.J.S. and B.S.M., and the Military Infectious Diseases Research Program (MI240008) grants. S.A.C. is a U.S. Federal employee. The work of this individual was prepared as part of official government duties. Title 17 U.S. C. §105 provides that “copyright protection under title is not available for any work of the United States Government.” Title 17 U.S.C. §101 defines U.S. Government work as work prepared by a military service member or employee of the U.S. Government as part of that person’s official duties. The views expressed are those of the authors and do not necessarily reflect the official policy or position of the Department of the Navy, Department of the Army, Department of Defense, or the U.S. Government. The following reagent was obtained through the NIH HIV Reagent Program, Division of AIDS, NIAID, NIH: Human Immunodeficiency Virus Type 1 HIV-1_Ba-L_ virus (ARP-510), contributed by Dr. Suzanne Gartner, Dr. Mikulas Popovic, and Dr. Robert Gallo. Anti-CD3 (OKT3) used in the current study was provided by the National Cancer Institute’s Biological Resources Branch Preclinical Repository. Some figures were created with BioRender. We thank Joshua Krueger for the preparation of plasmid reagents, Isabelle Finholm for her assistance with RNAscope, Dr. Geoffrey Hart for the kind gift of the P815 cells, and Dr. Alon Herschorn for the gift of the tet-inducible HIV-Envelope plasmid.

## Ethics Statement

All animal procedures reported herein were conducted under IACUC protocols approved by the Walter Reed Army Institute of Research/Naval Medical Research Command (WRAIR/NMRC) and the UMN (Protocol ID: 2401-41649A) in compliance with the Animal Welfare Act and by the principles outlined in the “Guide for the Care and Use of Laboratory Animals,” Institute of Laboratory Animals Resources, National Research Council, National Academy Press, 2011.

## Disclaimer

This material has been reviewed by the Walter Reed Army Institute of Research. There is no objection to its presentation and/or publication. The opinions or assertions contained herein are the private views of the author, and are not to be construed as official, or as reflecting true views of the Department of the Army or the Department of Defense.

## Author Contributions

L.K.T.: Data curation (lead), formal analysis (lead), investigation (equal), supervision (supporting), validation (lead), visualization (lead), writing-original draft (lead), writing-review and editing (lead).

P.P.: Data curation (supporting), formal analysis (supporting), investigation (equal), supervision (lead), project administration (equal), supervision (equal), validation (supporting), writing-review and editing (supporting).

J.W.C.: Data curation (supporting), investigation (equal), methodology(supporting), resources (supporting), validation (equal), review and editing (supporting).

M.E.C.: Data curation (supporting), investigation (equal), validation (supporting), writing-review and editing (supporting).

I.G.B: Data curation (supporting), investigation (supporting), validation (supporting), writing-review and editing (supporting).

K.L.B.: Data curation (supporting), investigation (supporting), validation (supporting), writing-review and editing (supporting).

Z.E.Q.: Data curation (supporting), investigation (supporting), validation (supporting), writing-review and editing (supporting).

A.F.K.: Investigation (supporting), methodology (supporting), resources (supporting), validation(supporting), writing-review and editing (supporting).

A.K.R.: Formal analysis (supporting), validation (supporting), writing-review and editing (supporting).

M.S.P.: Conceptualization (supporting), project administration (equal), resources (supporting), supervision (supporting), writing-review and editing (supporting).

S.A.C: Conceptualization (supporting), funding acquisition (supporting), methodology (equal), resources (equal), writing-review and editing (supporting).

V.V.: Conceptualization (supporting), funding acquisition (supporting), resources (equal), writing-review and editing (supporting).

B.S.M: Conceptualization (equal), funding acquisition (equal), project administration (equal), resources (equal), writing-review and editing (supporting).

P.J.S: Conceptualization (equal), funding acquisition (equal), project administration (equal), resources (equal), writing-review and editing (supporting).

