## Supplemental Information for "NK cell immunotherapy administered at the time of HIV recrudescence is associated with viral control"

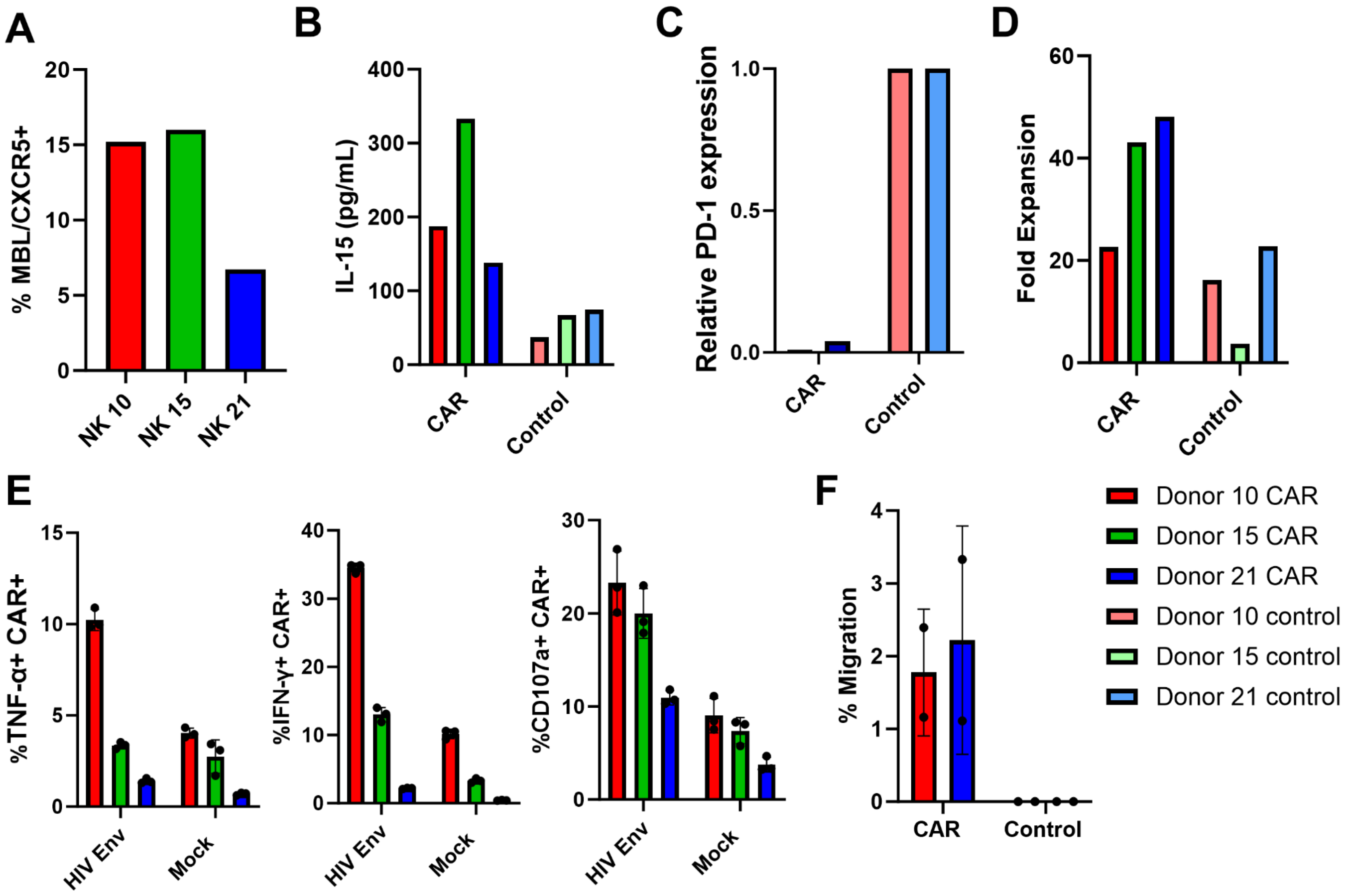


**Figure S1**: ***In vitro* testing of candidate NK infusion donors.** Three donors were engineered and evaluated for *in vitro* functionality: A) Expression of CAR (MBL) and CXCR5 molecules was confirmed in the cell product using flow cytometry. Cells were pre-gated sequentially on lymphocytes, singlets, live cells, CD56+, and CD3-. B) IL-15 expression in engineered NK cells was confirmed through IL-15 ELISA of cell culture supernatant and compared to unengineered control NK cells. C) PD-1 knockout was detected at the RNA level by RT-PCR with primers designed to detect the PD-1 sequence. D) Both CAR and control NK cells recovered from thaw and expanded at similar rates over the third expansion (days 12-19 overall). E) Pro-inflammatory cytokine (IFN-γ and TNF-α) and degranulation marker (CD107a) production was measured after CAR NK cells were cocultured with either HIV-Env or WT PBMCs. Measured via ICS assay. F) The percentage of cells that migrated to CXCL13 (ligand of CXCR5) was determined using a transwell plate migration assay. All assays were performed in duplicate or triplicate. In E and F, bars represent the mean and standard deviation. In all graphs, pink/red is donor 10, dark green/light green is donor 15, dark blue/light blue is donor 21.

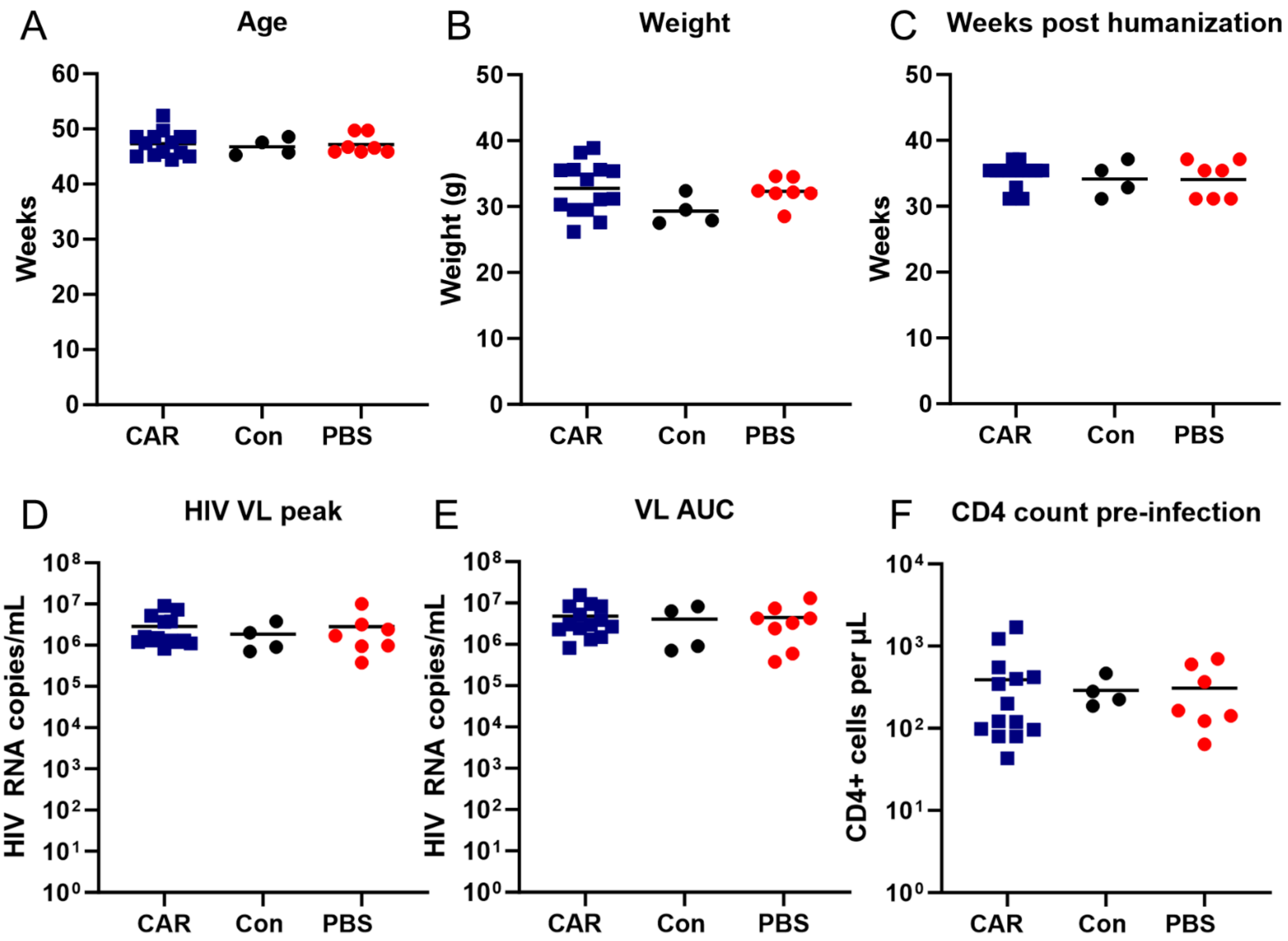


**Figure S2: DRAGA animal grouping information.** DRAGA mice were grouped for either CAR (blue), control (black), or PBS (red) treatment based on A) age at the time of treatment, B) weight at the time of treatment, C) amount of time since humanization D) HIV vRNA peak prior to ART treatment, E) Area under the curve of viral loads prior to ART treatment, and F) CD4 count prior to infection.

**Table S1: Summary of necropsy findings from animals with adverse outcomes during the DRAGA mouse study.**

| DRAGA Number | DRAGA Sex | Donor number and sex | Necropsy Time point | Symptoms |
|---|---|---|---|---|
| 28 | F | 1 (M) | 1 Week post-ART | >10% weight loss in one week<br>Anemia (pale blood)<br>Multifocal white spots on liver<br>Cyst on left ovary |
| 30 | F | 1 (M) | 4 Weeks post-infection | >10% weight loss in one week<br>Abnormal behavior: circling, lethargy |
| 32 | F | 1 (M) | 2 Weeks post-ART | >10% weight loss in one week<br>Anemia (pale blood)<br>Multifocal white spots on liver |
| 33 | F | 1 (M) | 20 Days post-treatment (PBS group) | >10% weight loss in one week<br>Abnormal behavior: circling, lethargy |
| 35 | F | 1 (M) | 4 weeks post-ART | Abnormal behavior: extreme lethargy, trouble standing<br>Anemia (pale ears and tail)<br>Splenomegaly with multifocal white spots<br>Hemorrhagic cyst on ovary taking up approx. 1/2 of the total abdominal cavity, blood and pus in surrounding tissue<br>Blood in cranial cavity |
| 42 | F | 2 (M) | 2 days post-ART interruption | Anemia (pale blood)<br>Blood in cranial cavity |

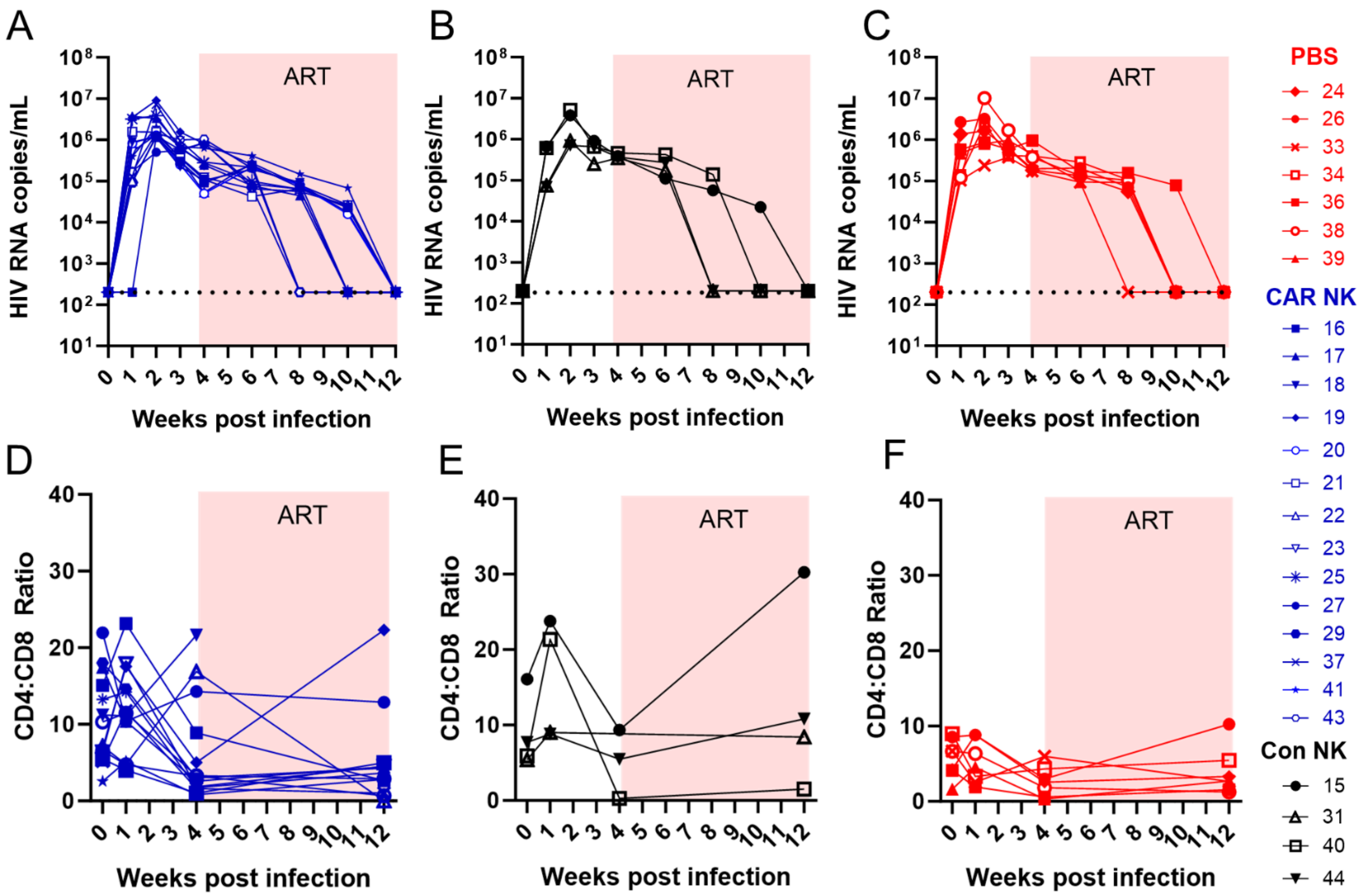


**Figure S3: Pre-treatment HIV viral loads and CD4:CD8 ratios.** Viral loads over time in A) CAR NK-treated animals (blue), B) control NK-treated animals (black), and C) PBS-treated animals (red). CD4:CD8 ratios over time in D) CAR NK-treated animals (blue), E) control NK-treated animals (black), and F) PBS-treated animals (red).

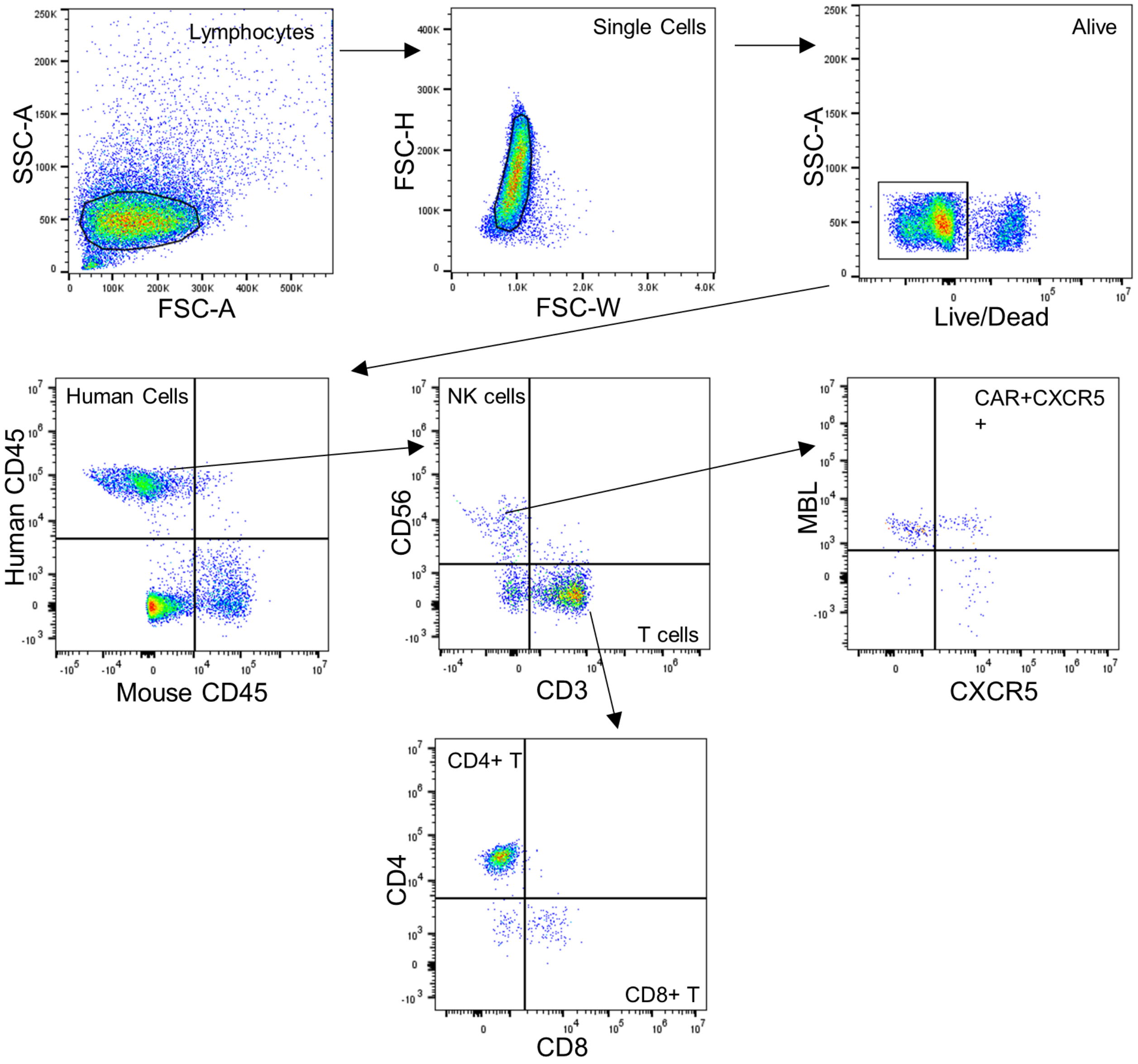


**Figure S4: Flow cytometry gating strategy for DRAGA mouse study.** Cells were gated Lymphocytes, singlets, alive cells, Human CD45+, Mouse CD45-. NK cells were gated CD56+, CD3-, and CAR NK cells were further identified by gating MBL+ CXCR5+. T cells were identified by gating CD3+ CD56-, and then CD4 or CD8+.

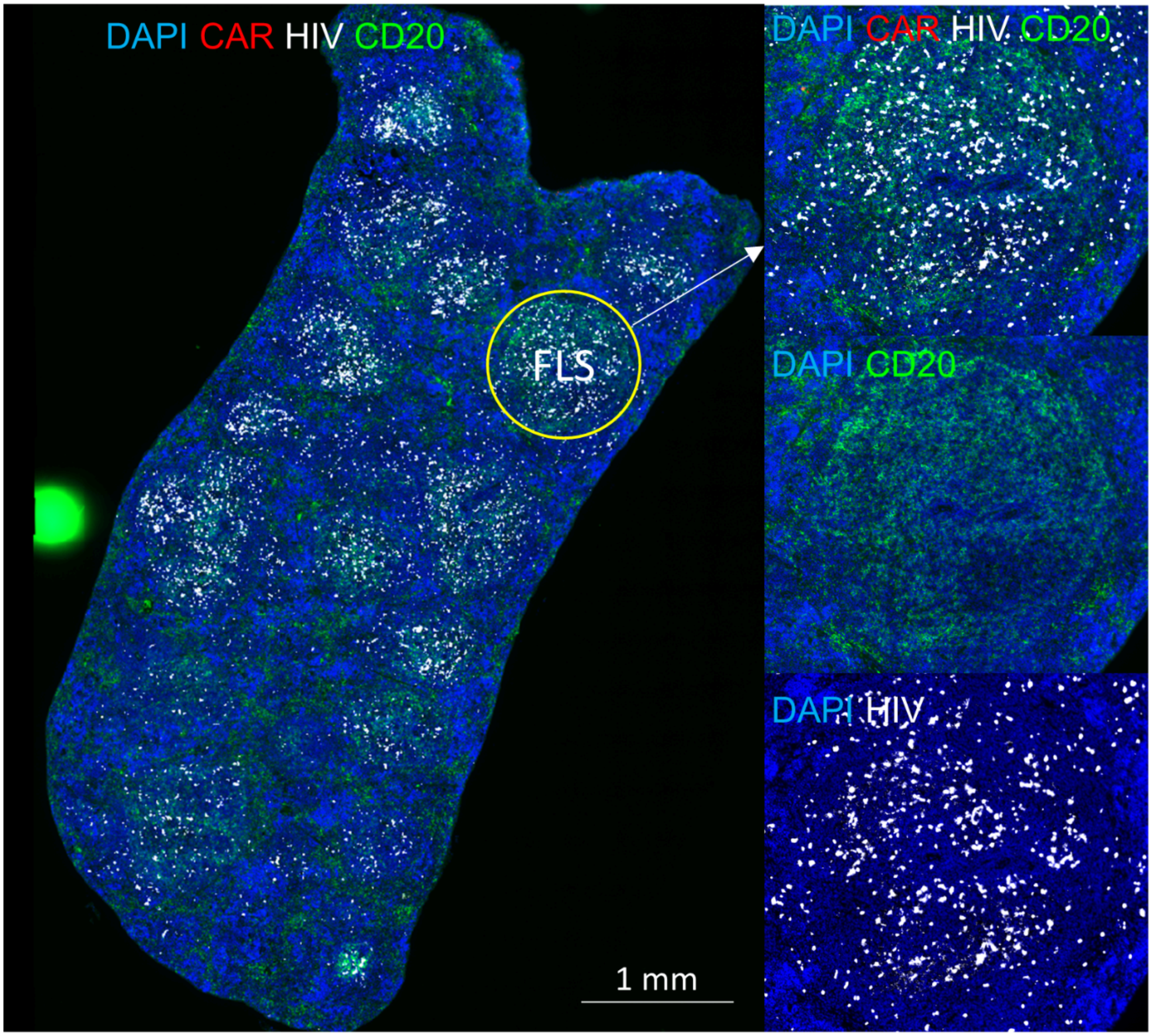


**Figure S5: HIV+ cell levels in the spleen at 6 DPT.** A) Spleen from DRAGA 25 at 6 DPT (necropsy) stained for DAPI (blue), CAR+ (red), HIV+(white), and CD20+ (green). FLS were identified by CD20+ morphology. On the right, enhanced images of the follicle with staining as indicated.

**Figure S6: Supporting Sequences**

CAR_P2A_CXCR5_T2A_IL-15:

MVRGVPFRHLLLVLQLALLPAATQGKKVVLGKKGDTVELTCTASQKKSIQFHWKNSNQIKILGNQGSF
LTKGPSKLNDRADSRRSLWDQGNFPLIIKNLKIEDSDTYICEVEDQKEEVQLLVFGLTANSDTHLLQGQS
LTLTLESPPGSSPSVQCRSPRGKNIQGGKTLSVSQLELQDSGTWTCTVLQNQKKVEFKIDIVVLAFQKAS
GGGGSKQVGNKFFLTNGEIMTFEKVKALCVKFQASVATPRNAAENGAIQNLIKEEAFLGITDEKTEGQF
VDLTGNRLTYTNWNEGEPNNAGSDEDCVLLLKNGQWNDVPCSTSHLAVCEFPIAAATTTPAPRPPTPAP
TIASQPLSLRPEACRPAAGGAVHTRGLDFACDIYIWAPLAGTCGVLLLSLVITLYCKRGRKKLLYIFKQPF
MRPVQTTQEEDGCSCRFPEEEEGGCELRVKFSRSADAPAYQQGQNQLYNELNLGRREEYDVLDKRRG
RDPEMGGKPRRKNPQEGLYNELQKDKMAEAYSEIGMKGERRRGKGHDGLYQGLSTATKDTYDALHM
QALPPRGSGATNFSLLKQAGDVEENPGPMNYPLTLEMDLENLEDLFWELDRLDNYNDTSLVENHLCPA
TEGPLMASFKAVFVPVAYSLIFLLGVIGNVLVLVILERHRQTRSSTETFLFHLAVADLLLVFILPFAVAEGS
VGWVLGTFLCKTVIALHKVNFYCSSLLLACIAVDRYLAIVHAVHAYRHRRLLSIHITCGTIWLVGFLLAL
PEILFAKVSQGHHNNSLPRCTFSQENQAETHAWFTSRFLYHVAGFLLPMLVMGWCYVGVVHRLRQAQ
RRPQRQKAVRVAILVTSIFFLCWSPYHIVIFLDTLARLKAVDNTCKLNGSLPVAITMCEFLGLAHCCLNP
MLYTFAGVKFRSDLSRLLTKLGCTGPASLCQLFPSWRRSSLSESENATSLTTFGSGEGRGSLLTCGDVEE
NPGPMRISKPHLRSISIQCYLCLLLNSHFLTEAGIHVFILGCFSAGLPKTEANWVNVISDLKKIEDLIQSM
HIDATLYTESDVHPSCKVTAMKCFLLELQVISLESGDASIHDTVENLIILANNSLSSNGNVTESGCKECEE
LEEKNIKEFLQSFVHIVQMFINTS
